# Enabling automated and reproducible spatially resolved transcriptomics at scale

**DOI:** 10.1101/2022.03.21.483636

**Authors:** Linnea Stenbeck, Fanny Taborsak-Lines, Stefania Giacomello

**Author notes:** Equal contribution (Linnea Stenbeck, Fanny Taborsak-Lines).

## Abstract

Tissue spatial information is an essential component to reach a holistic overview of gene expression mechanisms. The sequencing-based Spatial transcriptomics approach allows to spatially barcode the whole transcriptome of tissue sections using microarray glass slides. However, manual preparation of high-quality tissue sequencing libraries is time-consuming and subjected to technical variability. Here, we present an automated adaptation of the 10x Genomics Visium library construction on the widely used Agilent Bravo Liquid Handling Platform. Compared to the manual Visium library preparation, our automated approach reduces hands-on time by over 80% and provides higher throughput and robustness. Our automated Visium library preparation protocol provides a new strategy to standardize spatially resolved transcriptomics analysis of tissues at scale.

## Introduction

The advances made in RNA sequencing (RNA-seq) have revolutionized how we analyze gene expression, making it possible to study whole transcriptomes in a high-throughput manner without a priori knowledge^1^. With the introduction of single-cell RNA sequencing (scRNA-seq), an entirely new field of analysis was enabled^2^. The whole-transcriptome analysis of single cells has provided extensive insight into gene expression heterogeneity and cell-type composition^3^. Though powerful, both RNA-seq and scRNA-seq do not allow to retain the spatial information of tissues, which is essential for understanding cell-to-cell interactions and obtaining a more holistic comprehension of gene expression mechanisms.

To overcome this issue, several spatially resolved transcriptomics techniques have been developed. These can be divided into targeted and untargeted approaches. Targeted methods use specific probes for their genes of interest^4-7^ and are imaged through fluorescence, while untargeted approaches allow unbiased transcriptome-wide studies through next-generation sequencing^8-13^. One of these is Spatial Transcriptomics (ST)^10^, which combines histological analysis of tissue sections with the detection and visualization of their whole transcriptomes by attaching them to a glass slide coated with spatially barcoded poly-dT capture probes. Because of its widespread usage through the 10x Genomics Visium Spatial Gene Expression assay, robustness, reproducibility and throughput are becoming even more critical. One way to address these aspects is to automize the library preparation steps of the protocol^14-17^. In fact, it has previously been shown that the adaptation of sequencing library preparation on a robotic workstation increases not only the robustness and the throughput of the method^18,19^, but also reduces the labor. Moreover, an automated approach minimizes risks of cross-contamination and human error, as well as reduces costs by requiring less hands-on time.

Here, we present an automated protocol for the library construction of the Visium Spatial Gene Expression protocol using the Agilent Bravo Liquid Handling Platform, a widely used robotic workstation in genomics laboratories. Our approach increases the throughput and robustness of the library construction as well as reduces hands-on time, enabling large-scale efforts and potential applications in the clinic sector.

## Results

### Automation of the protocol

We developed the automation of the Visium Spatial Gene Expression protocol by dividing it into two parts. The first part is performed manually on the Visium Spatial Gene Expression slide, whereas the second is performed on the released material in an automated fashion (Figure 1A). The manual part is usually performed in one day (referred to as Day 1) and entails the Visium Spatial Gene Expression protocol^20^ from cryosectioning to size selection of the amplified cDNA material (Figure 1A). Briefly, tissue sections on the slide undergo fixation, staining, imaging, permeabilization, reverse transcription, second-strand synthesis and cDNA denaturation. After the cDNA denaturation step, samples are collected from the slide and analyzed by qPCR to determine the number of cycles for the cDNA amplification. Subsequently, samples undergo size selection and purification as well as a quality control using a capillary electrophoresis instrument, thus ending Day 1. On the second day (referred to as Day 2), the samples are transferred to a 96-well PCR plate together with the master mixes required for library construction and loaded onto the Agilent Bravo Liquid Handling Platform (hereafter referred to as robot).

**Figure 1:**
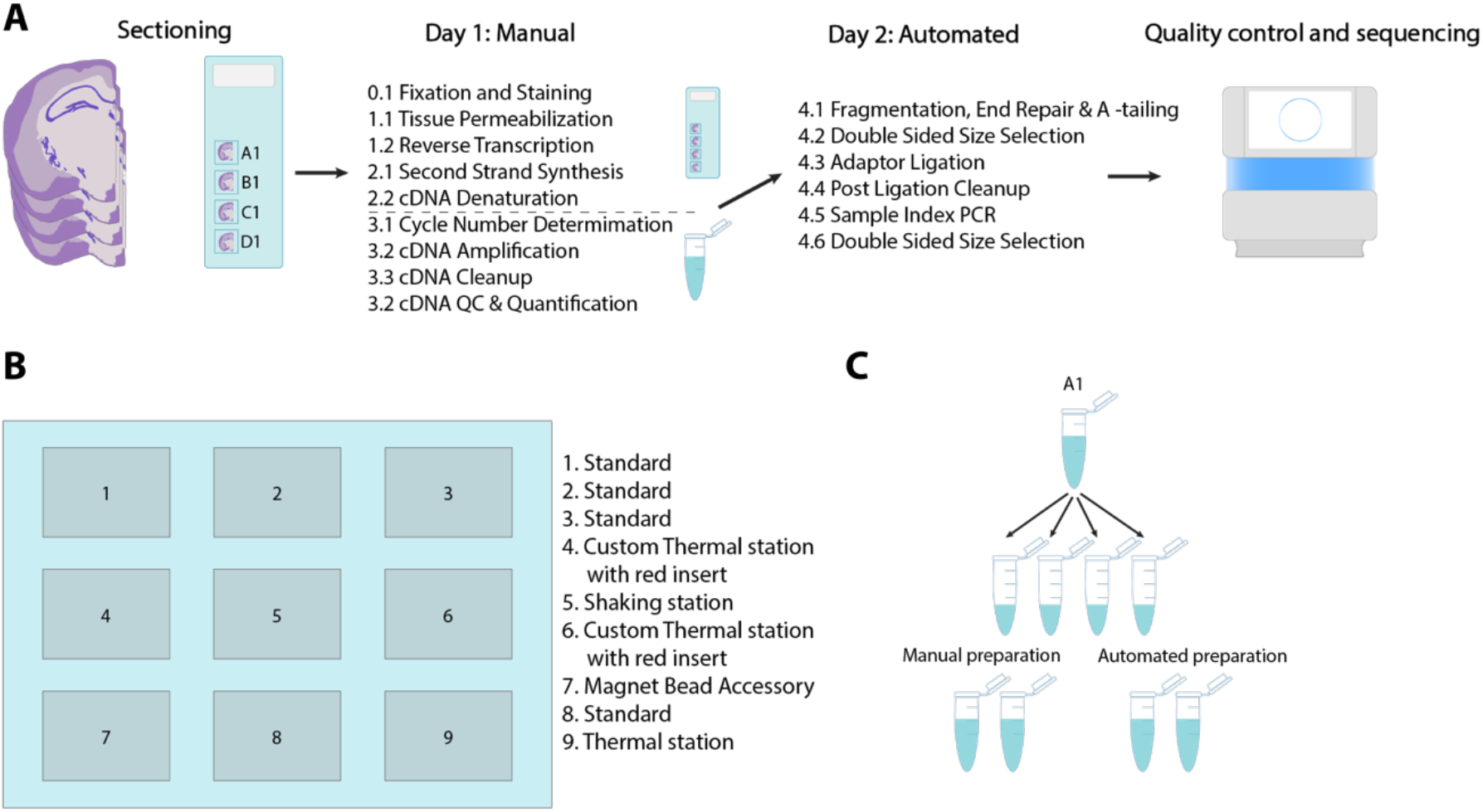
Overview of the protocol and setup. **A)** Schematic of the entire workflow; the protocol is divided into two parts, one manual and one automated. The manual part starts with tissue sections already sectioned and placed onto the Visium Gene Expression array. Day 1 (manual part) is performed on the array until cDNA denaturation, after which the material is released. The manual part ends with a quality control to ensure good libraries before continuing with the library construction. Day 2 (automated part) consists of library construction which begins with fragmentation and also ends with a quality control to determine library sizes. **B)** Layout of the working deck of the Agilent Bravo Liquid Handling Platform. Prior to starting the automated part, the working deck is loaded with a full tip box on position 2, an empty Nunc deep well 1.3 ml plate on position 3, a 2.2 ml deep well storage plate containing room temperature reagents on position 5, an empty 96-well PCR plate on position 6, an empty tip box on position 8 and a 96-well PCR plate containing the samples and reagents on position 9. **C)** Schematic of the division of samples to technical replicates for the comparison experiments. Well A1 of the Visium Spatial Gene Expression array is used as an example.

The working deck of the robot holds nine different positions, three of which are temperature controlled, one contains a shaking plate and one is a bead magnet accessory (Figure 1B). Apart from the reagent plate, the robot is also loaded with a full tip box, an empty tip box, an empty 96-well PCR plate to be used for incubations, an empty 1.3 ml Nunc deep well plate to be used for bead purifications, and a 2.2 ml deep well storage plate for room temperature (RT) reagents and waste.

Day 2 of the protocol begins by preparing the reagent plates, loading the robot with the previously mentioned objects, and starting the protocol. The robot performs the entire protocol without any manual intervention up to the preparation of the index library reaction, after which it pauses at 4°C to allow for sealing of the plate and its transfer to a thermocycler for the indexing PCR. Afterwards, the PCR plate is transferred back to the robot for the continuation of the protocol. Before the final elution step, the robot pauses at RT to allow for replacement of the PCR plate with a clean one. The final libraries undergo a post library construction quality control before sequencing by using a capillary electrophoresis instrument to determine the library size.

Our protocol allows for the preparation of up to 16 libraries simultaneously, with two different setups depending on the number of samples processed. The first setup allows to process up to 8 samples and the second one up to 16 samples, though the only difference is the preparation of the reagent plates. Our automated approach provides an 80% hands-on time reduction, from 125 min to 21 min (Table 1), despite a 20-minute overall increase compared to manual preparation. This is due to the added drying time during the bead purifications to ensure complete evaporation of the ethanol. Finally, to minimize manual intervention as well as plastic waste, we programmed the robot to reuse pipette tips throughout the entire protocol, when possible.

**Table 1:** Time comparison between the manual and automated library preparation. Times for the manual protocol are taken from the Visium Spatial Gene Expression protocol, with the reaction time subtracted from the total to get the hands-on time for the preparation of the samples. The hands-on time for the automated preparation is the preparation of the plates and the handling of the robot. The total time includes the manual handling as well as reaction time.

| Steps | Manual 1-8 samples |  | Automated 1-16 samples |  |
| --- | --- | --- | --- | --- |
|  | Hands on | Total | Hands on | Total |
| Thawing and setup | 5 min | 30 min | 15 min | 30 min |
| 4.1 Fragmentation, End Repair & A-tailing | 15 min | 50 min | - | 40 min |
| 4.2 Double Sided Size Selection | 30 min | 30 min | - | 43 min |
| 4.3 Adaptor Ligation | 10 min | 25 min | - | 17 min |
| 4.4 Post Ligation Cleanup | 20 min | 20 min | - | 33 min |
| 4.5 Sample Index PCR | 15 min | 40 min | 5 min | 40 min |
| 4.6 Double Sided Size Selection | 30 min | 30 min | 1 min | 43 min |
| Total: | 125 min | 225 min | 21 min | 246 min |

### Comparison between automated and manual preparations

To validate the automated protocol, we benchmarked it against the manual preparation of the Visium Spatial Gene Expression protocol. For this purpose, we selected two very different sample types: commercially available human reference RNA and mouse brain tissue. Human reference RNA was selected for its standardized nature, which minimizes batch variations and is therefore optimal for optimizing and comparing genomic experiments. Mouse brain tissue, on the other hand, is a well-characterized tissue type with distinct morphology, suitable to test the preservation of the spatial gene expression patterns processed through the automated protocol.

We considered four samples for the benchmarking analysis: two of them being human reference RNA as starting material, whereas the other two being mouse brain tissue sections (Supplementary Figure 1). We applied the Visium Spatial Gene Expression protocol Day 1 to all four samples. On Day 2, we divided each of the four samples into four technical replicates before the fragmentation, obtaining a total of 16 samples (Supplementary Figure 1). In order to have a direct comparison between the manual and automated preparation on the exact same material, we processed the four identical technical replicates deriving from the same original sample in pairs (one pair being two technical replicates processed manually, whereas the other one contained the two remaining replicates using the robot; Figure 1C). We analyzed the final library size profiles on a Bioanalyzer 2100 (Agilent) instrument and observed that the libraries prepared with the automated approach had a more equal distribution of the fragment sizes compared to those of the manually prepared libraries, suggesting a more robust procedure (Figure 2A).

**Figure 2:**
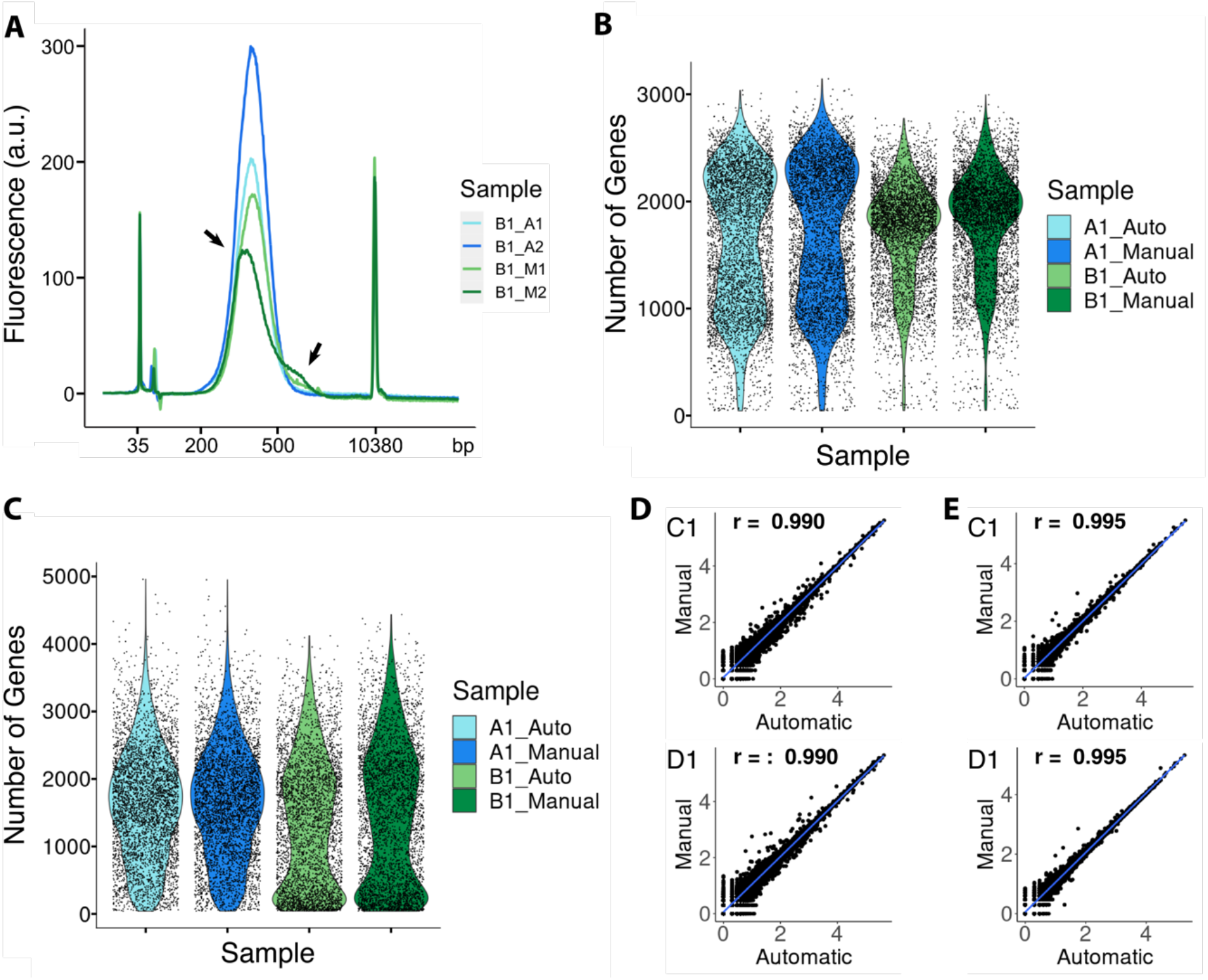
Evaluation of technical variability between manual and automated preparation. **A)** Evaluation of library sizes of all replicates from one of the tissue samples. Arrows indicate irregular distribution of fragment sizes for one of the manually prepared replicates of well B1. **B)** Distribution of gene counts per spot for each of the sequenced replicates for the samples originating from reference RNA. **C)** Distribution of gene counts per spot for each of the sequenced replicates for the samples originating from tissue. **D)** Pairwise correlations of the log10 normalized gene counts for the manually and automated prepared libraries between the replicates originating from reference RNA. **E)** Pairwise correlations of the log10 normalized gene counts for the manually and automated prepared libraries between the replicates originating from tissue.

Subsequently, to quantitatively compare the manual and automated library preparation, we sequenced one replicate from each preparation pair, sequencing a total of eight libraries (i.e., four derived from tissue section and four derived from reference RNA as input material, respectively). To conduct a fair comparison, we down-sampled each sample set (one set being the four samples originating from tissue and the other being the four samples originating from reference RNA) to the library with the least amount of reads in the corresponding set (60 million and 81.5 million, respectively) and analyzed all replicates separately. We observed very similar results between the reference RNA replicates, with an average number of genes per spot being 1635 and 1686 for the automatic and manual library preparation of the first sample, respectively, and 1712 and 1806 for the replicates of the second sample (Figure 2B). Moreover, we observed congruent numbers of genes per spot also between the manual and automatic library preparation in the two samples originating from mouse brain tissue sections. Specifically, we obtained 1622 and 1639 average genes per spot in the automatic and manual library preparation from the first mouse tissue section, respectively, and 1356 and 1488 for the other tissue section (Figure 2C). Overall, the average genes per spot within both sets of replicates were more similar than that between the samples prepared with the same approach (consecutive sections for the tissue samples), thus confirming that our automated approach was comparable to that of the manual protocol.

Next, we quantified the gene expression similarity between the automatic and manual library preparation per sample type and found a correlation of 0.990 between the libraries originating from reference RNA and 0.995 between the libraries originating from mouse brain tissue sections, respectively (Figure 2D-E). Lastly, we compared spatial gene expression patterns between the manually and automatically prepared mouse brain tissue sections. By visual inspection, major tissue domains, like for example cortex and hippocampus, presented highly overlapping spatial distributions of the number of genes per spot (Figure 3A). To confirm this observation, we performed dimensionality reduction and clustering of the gene expression captured in the manually- and automatically-prepared replicate independently. After matching the cluster identity between the replicates (Figure 3B), we calculated the number of spots per cluster in the two replicates and identified very similar values (Figure 3C). Finally, we visualized the spatial clusters and calculated the fraction of spots belonging to the same cluster between the two replicates, resulting in an overlap of 90% (Figure 3D). Taken together, these results confirm that our automated approach is highly comparable to the manual library construction in the Visium Spatial Gene Expression protocol.

**Figure 3:**
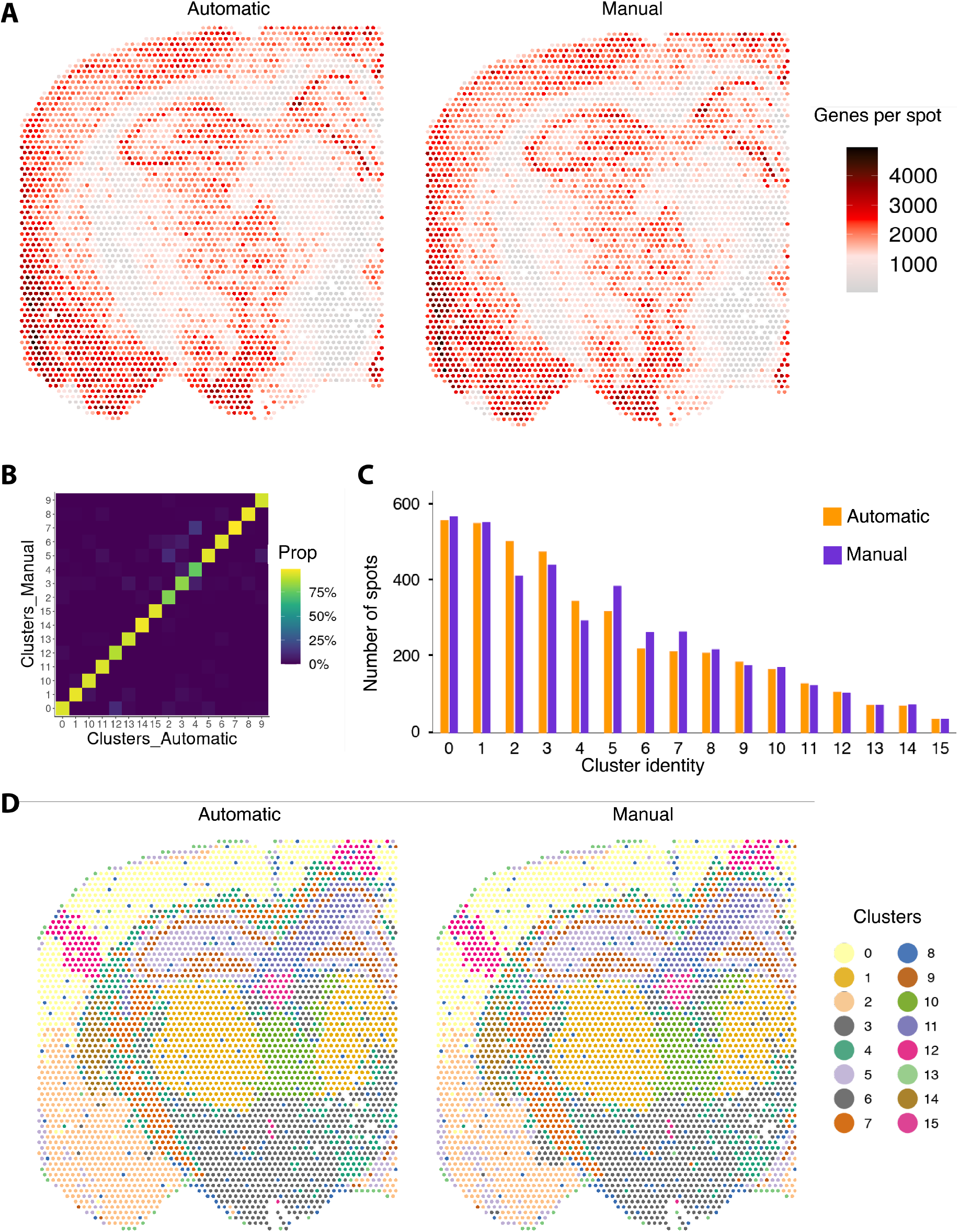
Evaluation of spatial variability between manual and automated preparation of sample A1 for mouse brain tissue. **A)** Spatial distribution of the number of genes in each spot between the two technical replicates of the same tissue section. **B)** Heatmap showing the percentage of spot overlap between clusters from the two replicates. **C)** Number of spots assigned to each cluster in both replicates. **D)** Spatial distribution of the clusters in the two replicates.

### Reproducibility testing

To evaluate the reproducibility of the automated protocol, we compared libraries obtained from the same starting material but prepared on three different robot runs. To this end, we used the remaining reference RNA and mouse brain tissue samples from the two different slides that were used for the comparison analysis, i.e., two of each sample type (Supplementary Figure 1). Before fragmentation, we divided these four samples into three replicates each and loaded them onto different positions within the 2 available columns of the reagent PCR plate across the three robot runs to remove batch effects due to the location (Supplementary Figure 1). In addition, we pooled reaction reagents across the three runs and applied both protocol setups for either up to 8 samples or up to 16 to ensure that the only variations that could occur were attributable to the robot.

After sequencing, we down-sampled each set of samples (one set being the six samples originating from reference RNA and the other one being the six samples originating from mouse brain tissue) to match the library with the least amount of reads in that set (57 million and 66.5 million, respectively) and normalized all samples separately. We observed very similar results between the reference RNA replicates, with an average number of genes per spot of ∼1340 for the first three replicates and of ∼1380 for the other three replicates (Figure 4A). Moreover, we found a high similarity trend between the mouse brain tissue section replicates, with an average number of genes per spot of ∼1500 for the first three replicates and of ∼1750 for the other three replicates (Figure 4B). To quantify the reproducibility, we calculated the gene expression correlation between all three replicates in the three different runs, obtaining very high Pearson correlation scores (Figure 4C-D), with an average value of 0.990 for the reference RNA samples and of 0.995 for the mouse brain tissue samples. Taken together, these results show that our automated approach provides a robust strategy for the Visium Spatial Gene Expression protocol with very high reproducibility.

**Figure 4:**
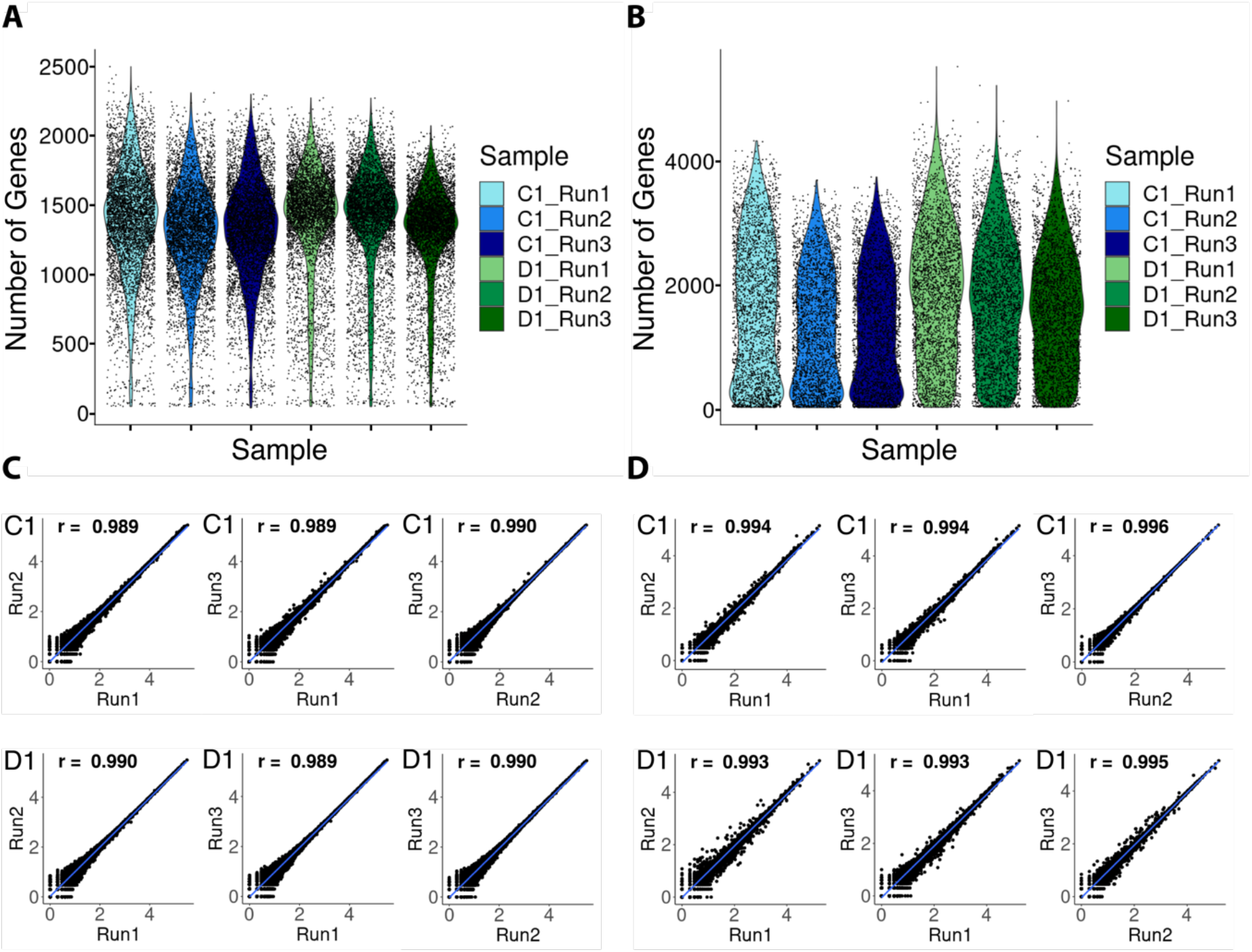
Evaluation of the reproducibility of the automated protocol. **A)** Distribution of gene counts per spot for each of the sequenced replicates for the samples originating from reference RNA. **B)** Distribution of gene counts per spot for each of the sequenced replicates for the samples originating from tissue. **C)** Pairwise correlations of the log10 normalized gene counts for the libraries prepared on different automated runs, from the replicates originating from reference RNA. **D)** Pairwise correlations of the log10 normalized gene counts for the libraries prepared on different automated runs, from the replicates originating from tissue.

## Discussion

The information provided through spatial gene expression analysis has proven essential to shed light on differences between similar cell types^21^. Among the commercially available methods that provide such information, the application of the 10x Genomics Visium Spatial Gene Expression assay is growing rapidly worldwide. As the usage of Visium is growing, so is the need for its standardization, especially considering that the protocol undergoes full manual execution. The manual execution of protocols, besides being time-consuming, causes the introduction of technical variation that can occur both between different individuals running the same protocol on the same sample and between different protocol days for the same individual. In contrast, an automated approach provides a decrease in these variations, increasing robustness and reproducibility as well as saving hands-on time.

As the method continuously increases in usage, several new applications could arise. Today, in the clinics, tissue analysis employs histopathology diagnostic tools, which are subjected to human interpretation^22^. One common histopathology tool is immunohistochemistry (IHC), which utilizes antibodies to visualize the distribution of specific antigens over a tissue^23,24^. IHC is a well-established method but is limited by throughput and human interpretation of the signal intensity of the targets of interest^25,26^. Having an approach that can quantify significant differences in gene expression levels in an unbiased manner would eliminate potential human errors, thus potentially enabling a digital pathology strategy. However, in order for the method to be integrated into the clinics, there is a need for more standardization to eliminate technical variations.

In this study, we have automated the Visium Spatial Gene Expression library construction on an Agilent Bravo Liquid Handling platform, a commonly present instrument in genomics facilities, allowing for robust library preparation for up to 16 samples simultaneously. Our automated protocol starts with the fragmentation of the amplified cDNA material, which is the first step of the library construction within the Visium Spatial Gene Expression procedure. It then proceeds with a fully automated execution with the exception of two instances where manual intervention is needed, one being the transfer of the plate to a thermocycler for the indexing reaction and the other being the change to a clean PCR plate for the final elution.

The automated approach reduces hands-on time by over 80% compared to the manual library construction, despite adding an extra 20 minutes compared to the manual preparation. This is mainly due to the bead drying time during all purification steps since there is no possibility to spin down the plate or make sure that all ethanol has evaporated before sample elution without pausing the robot and requiring manual intervention. Nevertheless, the amount of time saved in actual hands-on time is substantial, both in regards to the actual laboratory work and to the uninterrupted amount of time spent away from the experiment. When executing the manual protocol, the only time that could be spent away from the experiment is during incubations, which range between 15 and 40 minutes. Instead, our automated approach allows for 2.5 consecutive hours of hands-off time after starting the protocol. Moreover, it allows for the construction of 16 libraries simultaneously, doubling the number of samples that can be processed compared to the manual approach. Another important aspect of our automated approach is that it leads to highly comparable results with the manual approach and it is highly reproducible. Our results were obtained by analyzing two very different sample types such as human reference RNA and mouse brain tissue sections, thus confirming the robustness of our protocol.

In conclusion, we have created an automated approach for the library construction of the Visium Spatial Gene Expression protocol. This approach has proven to give a comparable outcome to the manual preparation, save hands-on time, provide scalability and reduce plastic waste by requiring fewer pipette tips compared to the manual preparation. Thus, we envision a widespread usage of our automated approach to the Visium Spatial Gene Expression protocol to enable more robust future studies, especially in the clinic sector.

## Methods

The Visium protocol was adapted on the Agilent Bravo Liquid Handling platform equipped with a Peltier thermal station with custom plate nest on positions 4 and 6, an orbital shaking station on position 5, a magnetic bead accessory on position 7, and a thermal station on position 9. It uses the same volumes as the standard Visium Spatial Gene Expression protocol but with an addition of 15µl Vapor-Lock (Qiagen, 981611) to each sample during the first incubation (fragmentation, end-repair and A-tailing) to prevent evaporation. The protocol allows for the preparation of up to 16 samples simultaneously, with the robot operating column-wise. There are two different setups depending on the number of samples used (one for 1-8 samples and one for 9-16 samples). The instructions, together with protocol files for the Agilent Bravo Liquid Handling Platform, can be found in the “Code Availability” section.

### Human Reference RNA preparation

Preparation of the universal human reference RNA (Agilent, 74000) was done according to the manufacturer’s manual. In short, the tube was centrifuged at 12000 x g for 15 min at 4°C, after which the supernatant was removed. The pellet was then washed in 70% ethanol before another centrifugation at 12000 x g for 15 min at 4°C was carried out. The pellet was then resuspended in 200 µl RNase free water and divided up into several 1:10 and 1:100 dilutions, resulting in tubes of reference RNA containing 1000ng/µl, 100ng/µl and 10ng/µl.

### Tissue handling, staining and imaging

The preparation of the mouse brain samples was performed following the Methanol Fixation, H&E Staining & Imaging for Visium Spatial Protocols (10x Genomics) ^27^. In short, four sections at 8 µm of fresh frozen mouse brain (Adlego) were placed on a Visium Spatial Gene Expression slide and stored overnight at -80°C. The slide was then transferred to 37°C for 1 min before being immersed and fixated in methanol (VWR EU, 20847.307) at -20°C for 30 min. After the fixation, the sections were dried by adding 500 µl isopropanol (Fisher Scientific, A461-1) for 1 min and then airdried. Next, the sections were stained with Mayer’s Hematoxylin (Agilent, S23309) for 7 min, the slide was then washed in Milli-Q water, and then the sections were incubated in 1 ml of bluing buffer (Agilent, CS702) for 2 min. After another round of washing in Milli-Q water, 1 ml of Eosin mix (Sigma-Aldrich, HT110216, 1:10 dilution in Tris-Acetic Acid Buffer) was added and incubated for 1 min before washing. The tissue sections were then dried for 5 min at 37°C and mounted with 85% glycerol (Merck, 104094) and a coverslip. Imaging was performed using the Metafer VSlide system at a magnification of 20x and the images were processed using the VSlide software. After imaging, the coverslip and remaining glycerol was washed off in Milli-Q water and the slide dried before transfer to the Slide Cassette.

### Library preparation before fragmentation

The preparation of the libraries was done following the Visium Spatial Gene Expression User Guide^20^ up to the Spatial Gene Expression Library Construction with the exception of the reference RNA, for which the protocol started with the reverse transcription and the reference RNA was added to the RT master mix. In short, permeabilization of the tissue sections was performed using the permeabilization enzyme (10x Genomics, PN2000214) for 20 min at 37°C, after which it was removed and each well washed with 0.1x SSC. Two reverse transcription mixes containing RT Reagent (10x Genomics, PN2000086), Template Switch Oligo (10x Genomics, PN3000228), Reducing Agent B (10x Genomics, PN2000087), RT Enzyme D (10x Genomics, PN2000216) and Nuclease-free water were prepared with one having 40 µl of Human reference RNA (100ng/µl) added and 40 µl of water removed. The mixes were added and the slides were incubated at 53°C for 45 min. After removal, 0.08M KOH was added to each well and incubated at room temperature (RT) for 5 min and then washed with EB (Qiagen, 19086). Next, a second strand mix containing Second Strand Reagent (10x Genomics, PN2000219), Second Strand Primer (10x Genomics, PN2000217) and Second Strand Enzyme (10x Genomics, PN200218), was added and incubated at 65°C for 15 min. After washing with EB, 35 µl 0.08M KOH was added to each well and incubated for 10 min in RT. Into each tube in an 8-tube strip, 5 µl Tris (1M, pH7.0) was added and then 35 µl of sample from the wells was added to a corresponding tube. cDNA amplification was carried out (16 cycles for the tissue samples and 14 cycles for the reference RNA) using a master mix containing Amp mix (10x Genomics, PN2000047) and cDNA primers (10x Genomics, PN2000089), the number of cycles was determined using a qPCR with the addition of KABA SYBR FAST (Sigma-Aldrich, KK4600) beforehand. The samples were then purified using SPRIselect (Beckman Coulter, B23318) at 0.6X and a Bioanalyzer 2100 (Agilent) was run as a quality control before storing the samples overnight at -20°C.

### Setup for the experiments

For both slides, the two first wells (A1 and B1) were separated into four individual replicate samples containing 10 µl each, where two were prepared using manual preparation and the other two using the automated approach. The other two wells (C1 and D1) for both slides were separated into three individual replicate samples containing 10 µl where each was run on different locations within the two columns and on separate runs on the robot, using both setups.

### Manual preparation

The four samples originating from tissue and the four samples originating from reference RNA were combined to the same strip-tube before continuing with the manual protocol. Fragmentation was carried out by incubating the samples in 32°C for 5 min and then 65°C for 30 min with EB and a fragmentation mix containing Fragmentation Buffer (10x Genomics, PN2000091) and Fragmentation Enzyme (10x Genomics, PN2000090). After fragmentation, a purification using SPRIselect at 0.6X and 0.8X was performed before continuing to the adaptor ligation. An adaptor ligation mix containing Ligation Buffer (10x Genomics, PN2000092), DNA Ligase (10x Genomics, PN220110) and Adaptor Oligos (10x Genomics, PN2000094) was added and incubated for 15 min at 20°C, after which a purification using SPRIselect at 0.8x was carried out. For indexing, Amp mix (10x Genomics, PN2000047) and an individual dual index (10x Genomics, PN3000431) were added to each sample and the indexing protocol was run in a thermocycler (13 cycles for the tissue and 14 cycles for the reference RNA), the number of cycles used for indexing was determined by the cDNA input calculated from the previous quality control. After indexing, a final purification was made using SPRIselect at 0.6x and 0.8x.

### Automated preparation

All reagents needed were taken out to thaw for 30 min during which the 2.2 ml deep well storage plate was prepared containing SPRIselect beads, Ethanol, EB and Vapor-Lock (Qiagen, 981611). Next, the master mixes were prepared according to the same protocol as the manual preparation and kept on ice. The reagent plate was then prepared containing the samples (the other four samples originating from tissue and the other four originating from reference RNA), the master mixes and the indexes. All plates were then loaded onto the robot according to the setup and the protocol was started. Before indexing, the protocol paused and the plate was sealed manually and placed in a thermocycler to carry out the indexing protocol, which was performed with the same number of cycles as their manual counterpart (13 cycles for the tissue samples and 14 cycles for the reference RNA). After indexing, the seal was removed and the plate placed back onto the robot. The protocol was then run until the last elution, for which another pause appeared in order to place a clean plate for the final elution. For the reproducibility testing, the same preparation was done using the same reagents but carried out at three different runs using different positions within the two usable columns.

### Time calculations

The times presented for the manual preparation are taken directly from the Visium Spatial Gene Expression protocol, where the reaction time is subtracted from the total in order to get the actual hands-on time for preparation of the samples. For the automated preparation, the hands-on time is timed during the experiment and includes the preparation of the plates and handling of the robot.

### Sequencing

After the library constructions, a quality control was performed on all samples using an Agilent Bioanalyzer High Sensitivity chip and the concentrations were determined by running a High Sensitivity Qubit assay (Thermo-Fisher, Q32851). For the comparison analysis, only one of the replicates (one from the automatic preparation and one from the manual preparation that originated from the same well) was sequenced, due to the similarities of the samples. The remaining 20 samples were then sequenced on the NextSeq2000 (Illumina) at a depth of around 65 million paired-end reads per sample. The forward read contained 28 nucleotides and the reverse read 150 nucleotides.

### Analysis

All libraries were down-sampled to the sample in the set with the least number of reads using Seqtk (https://github.com/lh3/seqtk.git) before analysis. Next, all libraries were pre-processed and mapped using Spaceranger v.1.2.1 (10x Genomics). The tissue libraries were mapped to the mouse genome (mm10 – 3.0.0) and the libraries with reference RNA were mapped to the human genome (GRCh38 – 3.0.0). All analysis was performed in R (v.4.0.4) using the STUtility package ^28^, the entire workflow can be reproduced and assessed in R markdown (see “Code Availability”). In short, all libraries were normalized individually using SCTransform and then merged to Seurat objects based on experiment (comparison or reproducibility) and based on well identity. Violin plots were generated using the function VlnPlot for each set of libraries. Next, Pearson correlation scores were calculated for each pairwise comparison for all the libraries and then plotted using ggplot. Dimensionality reduction and clustering were then performed on the tissue samples. Cluster identity was matched between the replicates by pairwise comparison between the clusters of both replicates and converting the cluster identity to match the cluster of the other replicate with the highest overlap. To ensure correct labeling we analyzed the proportion of spots overlapping between clusters. The percentage of spots with the same cluster identity between the replicates was calculated after correcting the cluster identities between the samples. Lastly, the clusters were plotted using ST.FeaturePlot.

## Data Availability

All raw reads for the libraries were deposited in the NCBI SRA under BioProject ID: PRJNA775889 (https://www.ncbi.nlm.nih.gov/bioproject/PRJNA775889). Output data from the 10x Spaceranger used for analysis was deposited on Mendeley Data at http://dx.doi.org/10.17632/3ngy9xvfwx.1. Data will be accessible upon publication.

## Code Availability

Code related to the analyses performed in this study as well as the protocol files for the Agilent Bravo Liquid Handling Platform can be found on GitHub at https://github.com/giacomellolab/VisiumAutomation. The code will be accessible upon publication.

## Acknowledgments

We wish to thank Lovisa Franzén for the help at the beginning of the project, as well as Sami Saarenpää, who also provided continued support throughout. We also would like to thank Yuvarani Masarapu and Ludvig Larsson for bioinformatics support. This study was supported by Formas, Vetenskapsrådet and Cancerfonden

## Author contributions

L.S. and S.G. designed the study and experiments. L.S. performed the experiments. F.T.L. created the automated protocol with input from L.S. L.S. analyzed the data with supervision from S.G. L.S. and S.G. wrote the manuscript with input from F.T.L.

## Supplementary Figures

**Supplementary Figure 1:**
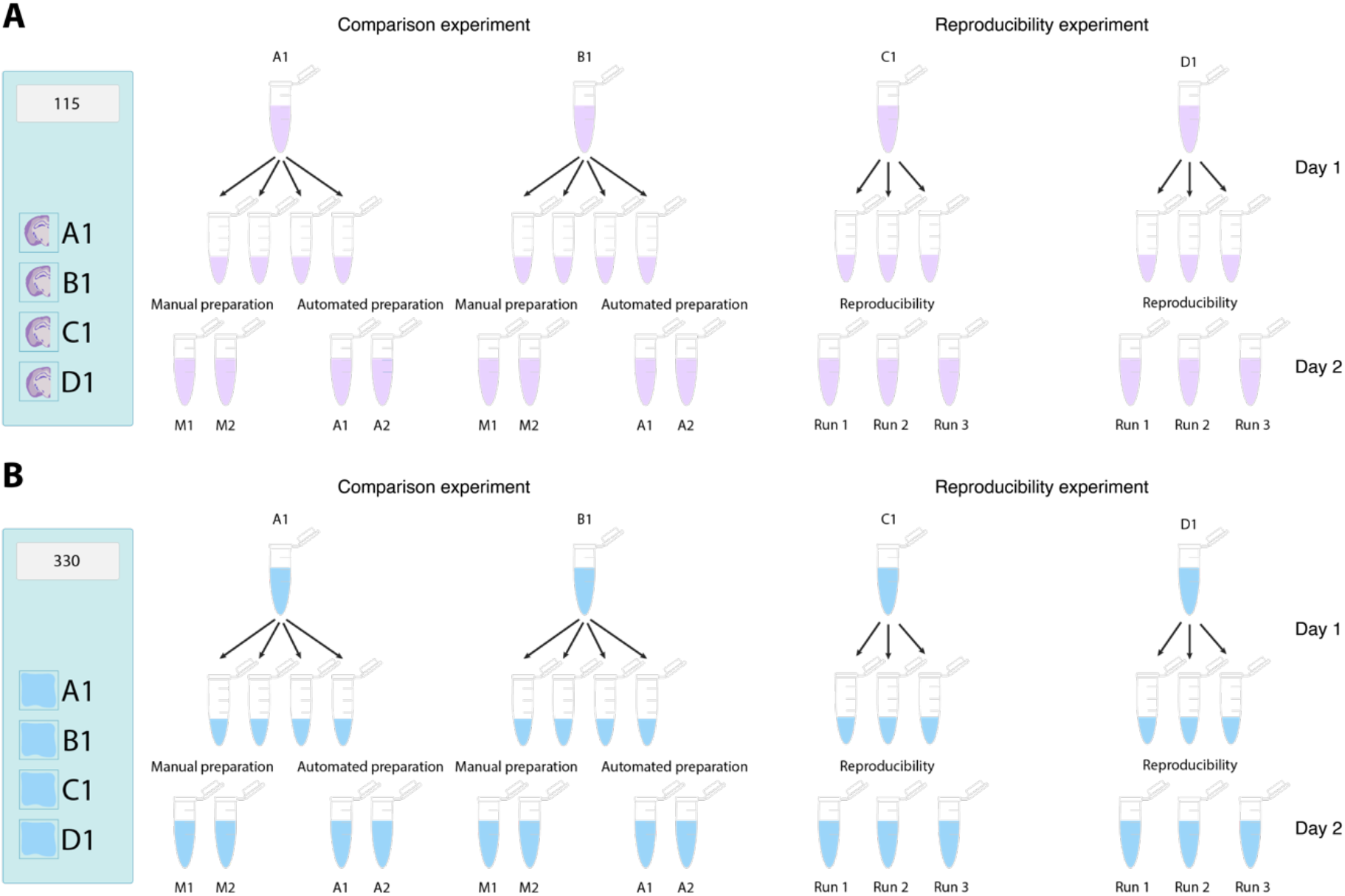
Overview of the division of samples to replicates for each experiment. **A)** Schematic of the division of tissue samples to replicates for the comparison experiment as well as the reproducibility experiment. **B)** Schematic of the division of reference RNA samples to replicates for the comparison experiment as well as the reproducibility experiment.

